# Rating enrichment items by group-housed laboratory mice in multiple binary choice tests using an RFID-based tracking system

**DOI:** 10.1101/2021.10.20.465117

**Authors:** Ute Hobbiesiefken, Birk Urmersbach, Anne Jaap, Kai Diederich, Lars Lewejohann

**Affiliations:** German Federal Institute for Risk Assessment (BfR), German Center for the Protection of Laboratory Animals (Bf3R), Max-Dohrn-Straße 8-10, D-10589 Berlin, Germany; Institute of Animal Welfare, Animal Behavior and Laboratory Animal Science, Freie Universität Berlin, Königsweg 67, D-14163 Berlin, Germany

**Keywords:** laboratory mouse, environmental enrichment, housing conditions, animal welfare, preference testing, C57BL/6J mice

## Abstract

There is growing evidence that enrichment of housing conditions of laboratory animals has positive effects on behavior, growth, and health. Laboratory mice spend most of their lives in their housing rather than in experimental apparatus, so improving housing conditions is a first-choice approach to improving their welfare. Despite the increasing popularity of enrichment, little is known about whether it is also perceived as being beneficial from the animal’s point of view. This is especially true due to the fact that ‘enrichment’ has become an umbrella term that encompasses a wide variety of different elements. Therefore, we categorized enrichment items according to their prospective use into the categories ‘structural’, ‘housing’, and ‘foraging’. In multiple binary choice tests we let 12 female C57BL/6J mice chose and rank 5 enrichment items per category. All possible pair combinations of enrichment items within each category were presented counterbalanced for a 46-hour period in a home cage based system consisting of two interconnected cages. A new analyzing method combined the binary decisions and ranked the enrichment items within each category by calculating worth values and consensus errors. Mice ranked the lattice ball (foraging), the rope (structural) and the second plane (structural) in upper positions. No clear preferences were determined for different types of housing enrichment during inactive times (light phase) whereas these objects were actively explored during the dark phase. Here the floorhouse and the paperhouse revealed high worth values. Overall, a high consensus error in ranking positions was observed reflecting strong individual differences in preferences. This highlights the importance of a varied enrichment approach as not all mice prefer the same item at all times. Given the known overall beneficial effects of enrichment, these data will help to provide appropriate enrichment elements to improve animal welfare and refine animal experimentation.

## Introduction

Attitudes toward animals as fellow living creatures have changed significantly in recent decades. There is growing concern about the conditions under which laboratory animals are kept, and it is therefore not surprising that legal requirements are also becoming increasingly demanding. In Europe, minimum requirements for housing laboratory animals are set out in EU Directive 2010/63^1^, which stipulates that animals must be housed according to the specific needs and characteristics of each species. Experimental animals should be provided with ‘space of sufficient complexity to allow expression of a wide range of normal behavior’. While the available space itself is a pressing issue for future improvements, the issue of complexity is usually approached through what is known as ‘enrichment of housing conditions’. It is reasonable to assume that additional enrichment opportunities in barren cages will create a more complex environment, which is likely to be appreciated by the animals^2,3^ and they are even willing to work for access to enrichment opportunities^4^.

However, it is important to note that ‘enrichment’ has become an umbrella term that encompasses a wide variety of different elements. Therefore, it must be kept in mind that by no means a uniformly accepted enrichment is meant when speaking of effects of enrichment ^2,3,5^. This being said, many research groups have indeed shown the benefits of enriched environments relative to conventional housing on well-being parameters in mice ^3,6^. Abnormal repetitive behavior expression, behavioral measures of anxiety, as well as growth and stress physiology were influenced positively by providing mice with a more varying environment using enrichment items ^7^. Access to enrichment lead to improved learning and memory function ^8,9^, increased hippocampal neurogenesis ^9,10^, attenuated stress responses and enhanced natural killer cell activity ^11^. Importantly studies showed no generalizable influence of a more diverse environment on variability of important parameters in biomedical research in mice ^12–14^. With regard to the workload of animal caretakers only a slight increase was noted while their overall assessment of providing enrichment in light of enhanced well-being for laboratory rodents was reported as good ^15^. Vice versa, there is increasing evidence, that keeping animals in standard housing conditions may be the negative factor in the development of behavioral disorders because of its impoverished character ^16^.

To create a more varied and stimulating environment, the size of the home cage can be enlarged, the group size increased, and stimulating elements can be provided ^17,18^. However, the human perspective does not necessarily reflect the wants and needs of mice^2^. Therefore, it is essential to ask the animals themselves about the adequacy of the enrichment items ^19,20^. To determine how different items are perceived by the animals themselves ^21^, animal centric strategies like preference tests will help to assess and rate different items ^20,22–24^.

From the three typically used preference testing designs ^23^, T-Maze, conditioned place preference, and home cage based preference tests, the last one seems to be the most appropriate for rating enrichment items. Especially when it comes to the avoidance of frequent animal handling and the opportunity to extend testing periods up to include a full circadian cycle or longer ^23^. Additionally, choice tests conducted within the home cage without the influence of an experimenter ^25,26^ correspond better to real laboratory keeping conditions. Home cage based testing systems usually consist of two ^27,28^ or more ^29,30^ connected cages with or without a center cage. In such tests mice are able to stay in their preferred surrounding and the cage that is chosen with the longer period of stay is regarded as the preferred one, or, in case of aversive properties, as the one least avoided ^23^.

For our preference test, we used the Mouse Position Surveillance System (MoPSS), a new test system designed and constructed in our laboratory ^31^ to ask for enrichment item preferences in female C57BL/6J mice, a widely used strain in biomedical research^32^. The MoPSS allows automatic long-term calculation of time spent in each of two interconnected cages for every individual mouse in a group. The determined dwelling time is used to conclude the choice between different enrichment items from the point of view of a mouse. The offered items were categorized and tested by their intended purpose of structuring the cage (structural enrichment), stimulating foraging engagement of the mice (foraging enrichment), and providing an alternative resting place (housing enrichment). To rank multiple items, we combined multiple binary choice tests and calculated worth values ^33^. In order to further evaluate the quality with regard to consistency of choice among individual mice and within groups of mice living in the same cage we used a recently developed method for analyzing worth value ratings ^34^. The overall aim of assigning worth values to specific enrichment items by comparison, is to provide scientifically based assistance for improving housing conditions of laboratory mice and thus increase animal welfare.

## Results

### Preference testing

The relative preferences (worth values, ranging from 0 to 1) and consensus errors (percentage of disagreement) of all 12 mice for the enrichment items of the categories foraging, structural, and housing during the entire 46-hours testing cycle and during active and inactive time are given in Figure 1.

**Figure 1.**
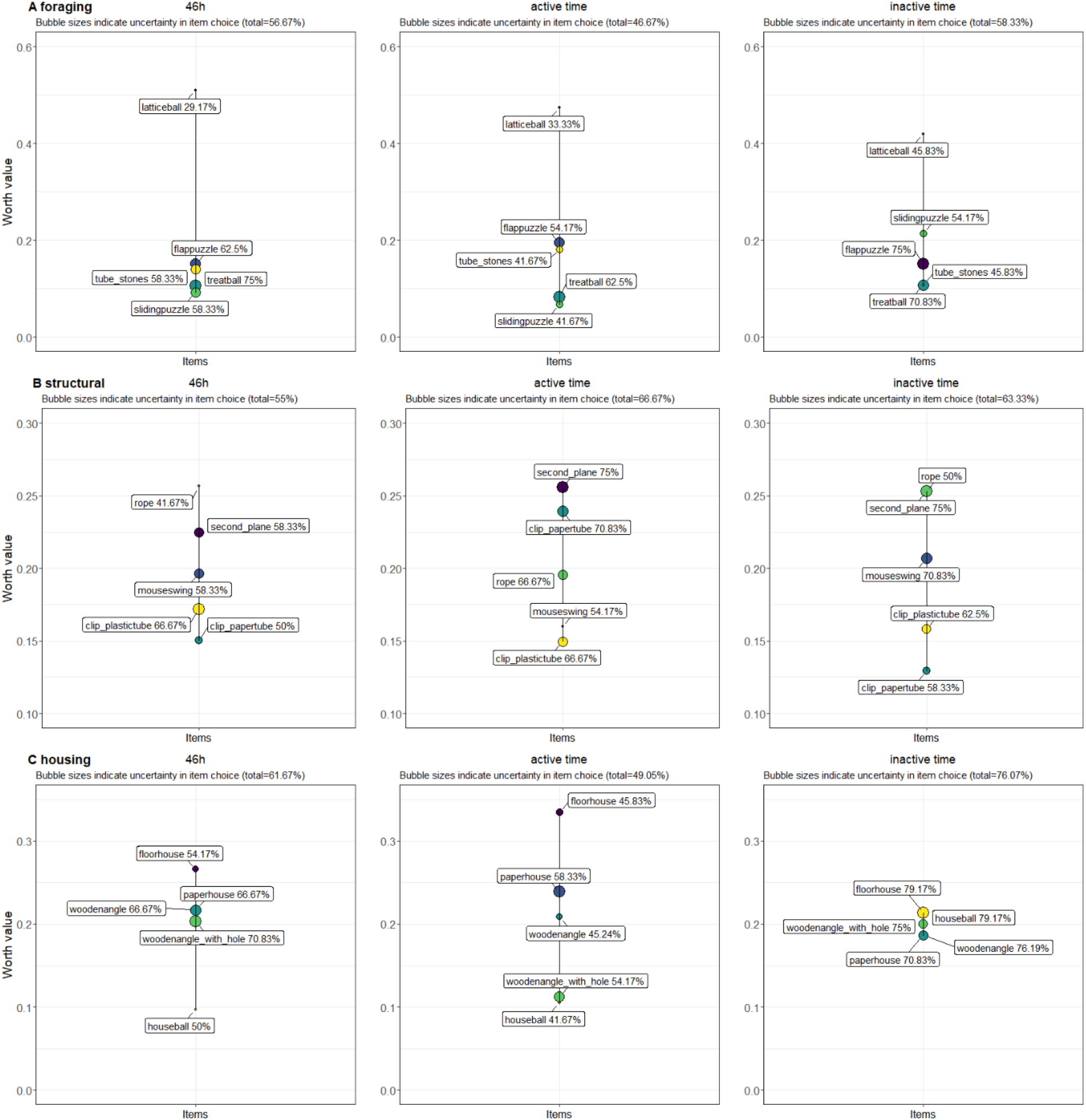
The relative preferences (worth values) and consensus errors (in percent) of all mice (n=12) for the tested enrichment items from the categories foraging, structural and housing in the single paired comparisons. The 46-hour period depicts the hole testing cycle whereas the active time depicts the dark phase of the testing cycle and the inactive time depicts the light phase of the testing cycle.

Mice preferred the lattice ball over all other *foraging enrichment* items during the 46-hour testing interval (mean worth value (WV): 0.51; consensus error (CE): 29.17 %), both during active (WV: 0.47; CE: 33.3 %) and inactive time (WV: 0.42; CE: 45.83 %).

Over the total time of 46 hours, the highest worth values regarding the *structural enrichments* were attributed to the rope (WV: 0.42; CE: 45.83 %). However, during the active time the second plane (WV: 0.42; CE: 45.83 %) was preferred while during the inactive time both, the second plane (WV: 0.25; CE: 75 %) and the rope (WV: 0.25; CE: 50 %) reached the highest worth values.

Out of the *housing enrichments* all mice preferred the floorhouse over 46 hours (WV: 0.27; CE: 45.83 %) and within the active time (WV: 0.34; CE: 45.83 %). During the inactive time the floorhouse (WV: 0.21; CE: 79.17 %) and the houseball (WV: 0.21; CE: 79.17 %) were equally ranked on first position.

Figure 2 illustrates the relative preferences (worth values) of the mice of *Group 1 (n=4), Group 2 (n=4)* and *Group 3 (n=4)* for the enrichment items of the categories foraging, structural and housing during the entire 46-hours testing cycle.

**Figure 2.**
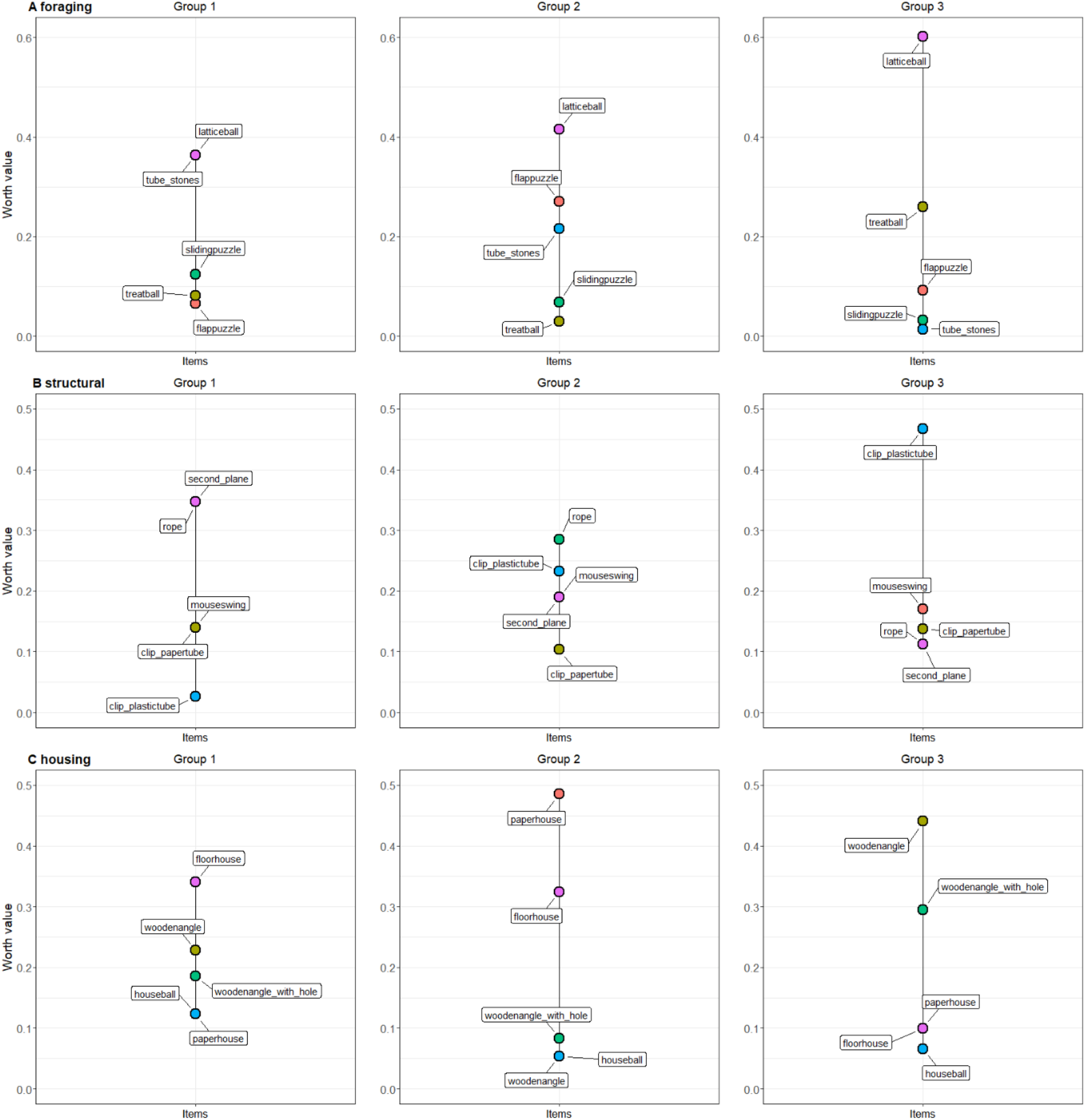
The relative preferences (worth values) of the mice from *Group 1* (n=4), *Group 2* (n=4) and *Group 3* (n=4) for the tested enrichment items from the categories foraging, structural and housing in the single paired comparisons over the entire 46-hour testing cycle.

Within the *foraging enrichments* group 1 ranked the lattice ball (WV: 0.36) and the tube with stones (WV: 0.36) on the first position, whereas group 2 and 3 ranked solely the latticeball (group 2 WV: 0.41; group 3 WV: 0.6) on the first position.

Among the *structural enrichments* group 1 ranked the rope (WV: 0.35) and the second plane (WV: 0.35) on the first position, group 2 ranked the rope (WV: 0.28) first and group 3 ranked the clip with the plastic tube (WV:0.47) first.

Analyzing the ranking positions of the *housing enrichments* on group level, group 1 ranked the floorhouse (WV: 0.34), group 2 the paperhouse (WV:0.49) and group 3 the wooden angle (WV:0.49) on the first position.

### Sample Size

Table 1 presents the results of the follow events, the influence events and the proportion of follow events and influence events of the transitions per mouse.

**Table 1.** The results from the follow and influence behavior analysis of 12 mice from the three experimental groups (1,2,3) of the complete data set.

| Group | Mouse | Transitions | Follow Events | Follow % | Influence Events | Influence % |
| --- | --- | --- | --- | --- | --- | --- |
| 1 | 1 | 11608 | 69 | 0.594 | 68 | 0.586 |
| 1 | 2 | 11132 | 54 | 0.485 | 73 | 0.656 |
| 1 | 3 | 10919 | 77 | 0.705 | 58 | 0.531 |
| 1 | 4 | 9955 | 58 | 0.583 | 59 | 0.593 |
| 2 | 1 | 10387 | 229 | 2.205 | 183 | 1.762 |
| 2 | 2 | 10224 | 199 | 1.946 | 193 | 1.888 |
| 2 | 3 | 8442 | 207 | 2.452 | 213 | 2.523 |
| 2 | 4 | 7557 | 143 | 1.892 | 189 | 2.501 |
| 3 | 1 | 7480 | 93 | 1.243 | 81 | 1.083 |
| 3 | 2 | 8440 | 97 | 1.149 | 96 | 1.137 |
| 3 | 3 | 7428 | 88 | 1.185 | 89 | 1.198 |
| 3 | 4 | 6013 | 64 | 1.064 | 76 | 1.264 |

The mean proportion of follow events in the transitions was 1.39% and the proportion of influence events in the transitions was 1.31%. If the follow interval was increased to 3 s, the proportion of follow events increased to 4.73 %.

Figure 3 depicts the ratio of follow and influence rate for all mice. Six mice showed very similar numbers of influence and follow events. Accordingly, they are close to the dividing line with the straight line slope of 1. Whereas the other six mice diverged from the dividing line either towards higher influence ratio or higher follow ratio. The proportion of follow events was lowest in the mice of group 1 and highest in group 2. Overall, the three groups appear to cluster, with animal within the same cage showing more similar scores than animals from different cages.

**Figure 3.**
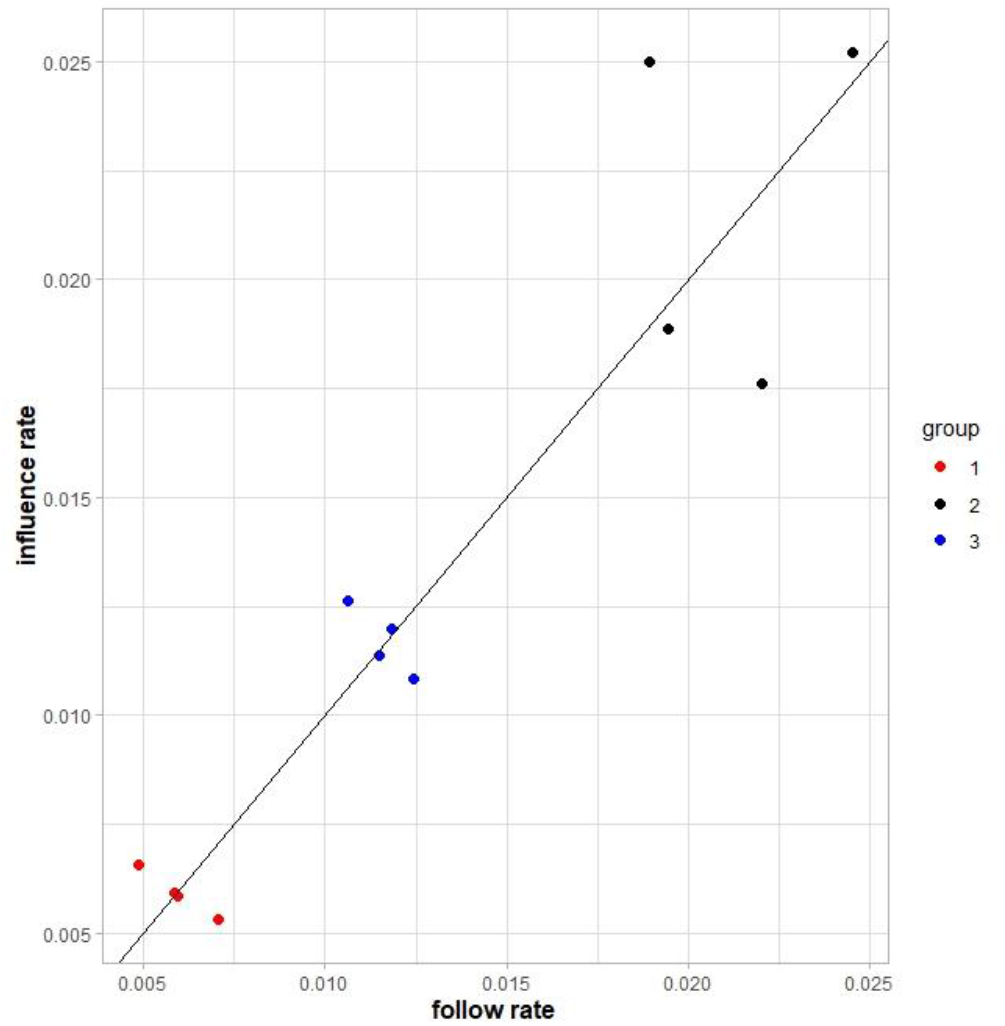
The ratio of the follow rate and influence rate from the follow and influence behavior analysis of 12 mice from the three experimental groups (1,2,3) of the complete data set.

## Discussion

The aim of this study was the evaluation of enrichment elements from the perspective of group housed female C57BL/6J mice. In a series of home cage based binary preference tests, mice could choose between different enrichment elements. The combined data were used to rank the items according to their worth value and to calculate the degree of disagreement in item selection between mice measured as consensus error (CE).

All choice tests were performed while the mice were in their respective social group in one out of three cages with four mice each. We conducted an analysis of follow and influence behavior, which shows how attached individual choice is to decisions of conspecifics. Data revealed that the three groups indeed did not come to the same conclusion with regard to choosing preferred items. However, there was no considerable attraction to individual mice that could explain the respective preference as a trend triggered by individual influencer mice. Overall, a mean follow rate of 1.39 % is reflecting a negligible direct impact on individual choices. Even if a more conservative follow interval was applied, more than 95% of all cage changes were not directly related to an influencer. Thereby we could demonstrate that group housed mice can explore a choice test apparatus without being directly led by others and thus an independent choice is likely. Nevertheless, testing groups of mice will remain a challenging issue with regard to choosing the correct statistical unit ^31,35,36^. Especially during the inactive phase, the location of a shared nest might influence the choice of the group. Mice are social animals in nature, and in accordance to underlying legislation^1^, single housing should be avoided under experimental conditions if possible. Furthermore, it is arguable whether choice decisions of individually kept animals can be transferred one-to-one to animals in a social group resembling realistic laboratory conditions ^31^. Thus, we decided against testing individual animals and used the option of the home cage based choice experiment to study the mice as socially living animals within the group ^37^. Furthermore, in addition to analyzing the results of all mice over the total test duration of 46 h, we subdivided the results into an active phase (dark) and inactive phase (light) of the mice ^4,38^. This served to evaluate possible preferences associated with active (e.g., climbing, gnawing) or inactive (e.g., sleeping, resting) behaviors of the enrichment items by the mice.

To investigate whether the mice agreed in their choice of preference we calculated a consensus error to display the amount of disagreement. Low scores indicated a high agreement, whereas high scores reflect a low agreement. Evaluation at the level of all mice of the three groups revealed a high average consensus error in all analyses and thus a lower agreement in choice, indicating different perceptions of enrichment within a group of mice. The individual group analysis showed that the rank positions of the tested enrichment elements varied greatly within their categories, which resulted in a high consensus error in total. Our assessment of follow and influence rates showed that this cannot be explained by dominance and following behavior. Therefore, the social dynamics underlying choice within a group are deemed to be more complex. In addition, the test items were freely available through the preference test, so the mice may not have perceived the test as forcing them to choose one or the other. This consideration is probably of greater importance if the difference in attractiveness between the objects is not very large. Indeed, it the CE is larger in rankings with low valence ranges compared to large valence ranges in the data provided with the R-package SimsalRbim ^34^.

### Foraging enrichments

were ranked with closely spaced worth values in all assessments. Only the lattice ball stands out with a high worth value, both at the group level and at the overall level. This is also reflected in the consensus error, which was the lowest in all calculations for the entire period at 29.17 % (CE in 46 h of all mice). Unlike the other enrichment items in the same category, the lattice ball was attached to the cage top using a metal ball chain, while the tube with stones, the flappuzzle, the slidingpuzzle, and the treatball were placed on the floor, resulting in high visual and functional differences. Due to the fact that after pulling paper out of the ball and eating the millet, the mice were still able to interact with the ball as a moving object to gnaw at or to climb on, it might have been more interesting and hence preferred to interact with. Moreover, mice used the pulled-out paper strips from the grid ball as additional nesting material (data not shown, observation during cleaning process) and combining different materials for nestbuilding has already been found to be common in mice ^39,40^. Indeed, nesting material is highly valued by mice^41,42^ and the motivation to build nests is strong^43,44^. The direct association of the lattice ball with supplementary nesting material may explain the preference for this side of the testing system also during the inactive phase of the mice. Our parallel study investigating the use of the lattice ball in the home cage showed that the active interaction with this design element was less frequent than with other foraging elements of the same category during the active phase (Hobbiesiefken et al., subm. ^45^). However, in that study the use was evaluated by direct observation during a 30-minute period in the presence of other enrichment elements from other categories. In the present study the data was obtained over two circadian cycles in a binary choice test which might be more conclusive with regard to the overall attractiveness. The elements ‘Treatball’, ‘slidingpuzzle’, ‘flappuzzle’, and the ‘tube filled with stones’ led to inconclusive worth values and thus low ranking positions in the evaluation at individual group and overall level. Nevertheless, they might serve as cognitive stimulation for mice and enable natural behaviors like burrowing and foraging. This is especially true when considering the high active usage while the elements were filled with millet seeds as additional treats.

### Structural elements

did also not reveal a complete unequivocally ranking which is again indicated by high values for the consensus error. However, within this category the second plane and the rope were highly ranked during both the active and inactive time of the mice. The second plane serves as a climbing element as well as for gnawing and as a refuge and sleeping place. The multifunctionality offers a wider range of possibilities for interaction compared to simpler climbing enrichments (i.e., mouseswing, clip with paper or plastic tube, rope). This is supported by a high rate of use of the second plane which we found in a comparative behavioral analysis (Hobbiesiefken et al., subm. ^45^). Leach et al.^46^ also acknowledged a platform-like insert for mouse cages as an appealing enrichment element for mice with its dual function as a resting place and as an object that encourages exploration, jumping, and hiding. In addition, we observed that mice frequently built their nests under the second plane, both, during the previous housing period as well as under the test conditions. The other structural enrichment, also preferred, was the rope. However, the evaluation of short-term usage in a previous study revealed this item, along with other climbing enrichments that were fixed at the cage top, to be less used when it was presented in a combination of enrichments (Hobbiesiefken et a., subm. ^45^). The rope was made of hemp and similar to the paper strips derived from the lattice ball, fragments of gnawed hemp ropes were used as additional nesting material. Therefore, the known attractiveness of nesting material ^41,42^ and the strong motivation to build nests^43,44^ might explain the high rank of the rope. This again shows that long-term observations are helpful to obtain more conclusive information about the overall attractiveness of the respective enrichment elements. Gjendal et al.^47^ found hemp ropes to strengthen the participation in social behavior and encouraging climbing and gnawing behavior in male mice without adverse effects on anxiety levels, stress and aggressive behavior. Hemp ropes can therefore be applied as a simple and inexpensive enrichment for mice and serve as climbing, gnawing, and supplemental nesting material.

### Housing enrichment

worth values were closely spaced, with apparent differences between groups, and elements partially achieving a reversed ranking. Accordingly, the consensus errors were considerably high for the overall rating of housing enrichment. Interestingly, van Loo et al. ^30^ found a comparable paperhousing to be preferred over a triangular plastic house. Therefore, we expected the paperhouse to be valued highly more consistently, however, this could not be confirmed unequivocally in our preference tests. Indeed, the paperhouse achieved the first ranking place in group 2 but was amongst the last ranking positions in group 1 and 3. Nonetheless, the paperhouse achieved the second place rank during the total and during the active time in the overall ranking. Indeed, a video observation revealed the frequent use of the paperhouse during the active phase of the mice (Hobbiesiefken et al., subm. ^45^). Apparently, the light weight and easily manipulated structure makes the paperhouse attractive as a movable and changeable object with which the home cage can be actively configured. The floorhouse was also rated highly in the active phase and seems to promote behaviors such as climbing, hiding and exploring more strongly. Due to its platform-like structure, it also offers a larger surface area for these types of behavior. Conversely, the houseball provides the least surface and was ranked to the lowest positions. During the inactive phase, no housing enrichment achieved a clear preference and all houses ranked closely spaced. This supports the hypothesis that nest boxes are also perceived as important exploration objects for mice rather than a mere refuge and sleeping area ^42,48^. This also shows that when mice are asked about their preference for provided items, the answer may be based on a different way of using these items than was expected by the experimenter. In general, mice prefer a cage with a nest box to a cage without a nest box^48^. Provision of nest boxes and nesting material increases animal welfare without negatively impacting data variability ^15^. Therefore, nesting material and nest boxes should be provided as a standard enrichment in mice ^3^. Since the nest box serves more than just shelter, the choice of design should also take into account the activity-promoting effect of the housing enrichment. Therefore, factors such as additional space or the changeable structure make the floorhouse and paperhouse recommended.

To determine the effectiveness of enrichment items, it is essential not only to conduct preference tests, but also to examine the ways in which enrichment items are used ^21^. Evaluation of the type and amount of interaction via behavioral analysis is therefore deemed an important component to create more species-appropriate housing conditions for mice (Hobbiesiefken et al., subm.^45^). Although we cannot provide a statement about the motivational strength^4,23,49^, the experimental design used here allows ranking of the different design elements. Determination of motivational strength can be achieved through consumer demand tests and represents the price an individual is willing to pay for access to certain enrichment elements ^4,20,24^. Nevertheless, our study shows that when mice have a say, judgments about a reasonable type of enrichment can be made in a somewhat more fact-based manner.

Overall, the high consensus errors in our study, especially for housing and structural enrichment, reflects individual differences in the assessment of the different enrichment elements from the perspective of each mouse. It should also be borne in mind that objects that are very similar cannot always be clearly distinguished from each other in terms of their valence ^34^. However, the fact that not all animals have always made a clear choice does not in any way indicate in principle that enrichment is superfluous. On the contrary, a comprehensive body of literature ^2,3,14–16,21,4,6–11,13^ shows positive effects of enrichment. From our study, in addition to practical recommendations, we can also derive the possibility of using different enrichment elements as a means of variation.

Indeed, to create an interesting and stimulating living environment for mice, it is important to provide variety through regular exchanges. Varied housing can help prevent behavioral deprivation^50,51^ and behavioral disorders^52^ in laboratory animals by enabling species-specific behaviors. Furthermore boredom in laboratory animals^53,54^ in its severe chronic forms shares symptoms with learned helplessness and depression and should therefore be treated as an important animal welfare concern^55^. It should be the ambition of every experimenter to improve the well-being of laboratory animals and thus enhance the quality of animal experiments^56^. Legal husbandry regulations^1^ should indeed be considered as a minimum requirement that does not place an upper limit on the genuine improvement of living conditions of laboratory animals ^22^.

## Conclusions

In our study, preferences for different enrichment items were elicited in female C57BL/6J mice using a home cage based preference test system. This easy-to-use method for obtaining information on worth values for different enrichment items for mice facilitates decisions on the use of enrichment in laboratory husbandry. We show that mice discriminate between different enrichment items, although not all animals always agree.

As foraging enrichment, the lattice ball with its multifunctional character of activity stimulation and its content of paper strips as additional nesting material achieved high worth values. A rope made of hemp is also highly valued as a structural element for climbing, gnawing, and providing additional nesting material. A wooden second plane was also favored as a structural enrichment, being used both as a resting place and for active engagement. No clear preferences were found for the type of housing during the inactive period of the mice. However, the houses serve more than just for sleeping, so structure-creating as well as manipulable houses are rated as advantageous.

High consensus errors within the studied rankings suggest a strong individuality in the perception of the enrichment elements. Therefore, a multifaceted enrichment approach should be considered to meet the needs of individual mice. Variation of enrichment elements also serves as a countermeasure to boredom, which can easily develop in small monotonous housing environments and lead to behavioral abnormalities. Increasing the complexity of housing for laboratory mice toward a more stimulating environment allows them to exhibit a more species-specific behavioral repertoire, potentially leading to more reliable animal models in biomedical research.

## Materials and Methods

### Ethical approval

All experiments were approved by the Berlin state authority, Landesamt für Gesundheit und Soziales, under license No. G 0069/18 and were in accordance with the German Animal Protection Law (TierSchG, TierSchVersV). The study was pre-registered in the Animal Study Registry (ASR, DOI 10.17590/asr.0000162).

### Animals and housing condition

Twelve female C57BL/6J mice were purchased from Charles River Laboratories, Research Models and Services, Germany GmbH (Sulzfeld). The sample size was chosen to ensure a statistical power of 80 % and an alpha value of 0.05. Due to the exploratory experimental approach, the effect size is unknown and had to be estimated on the basis of published studies with comparable experimental designs as well as own experiments from our laboratory. The mice were 7-8 weeks of age upon arrival in the animal facilities. Mice were randomly allocated to groups of four animals in Makrolon type III cages by a researcher not involved in the experiment; animals were alternately assigned to the groups (1,2,3) to avoid bias. During the first three weeks the animals were housed in groups of four animals in type III Makrolon cages (L × W × H: 425 × 265 × 150 mm, Tecniplast, Italy) with aspen bedding material (Polar Granulate 2-3 MM, Altromin), paper (cellulose paper unbleached 20×20 cm, Lohmann & Rauscher International GmbH & CO KG) and cotton roll nesting material (dental cotton roll size 3, MED-COMFORT), a 15 cm transparent plexiglas tube (Ø 4cm PMMA xt®, Gehr®) and a red triangle plastic house (mouse house, TECNIPLAST®). They were provided with regular rodent food (autoclaved pellet diet, LAS QCDiet, Rod 16, Lasvendi, Germany) and tap water *ad libitum*. Room temperature was maintained at 22 °C (+/- 2), room humidity at 55 % (+/- 15) and a 12/12 light/dark cycle regimen (lights off 20:00) with simulated sunrise between 7:30 and 8:00 using a Wake-up light (HF3510, Philips, Germany). To further implement refinement procedures according to the 3Rs ^57^ all mice were trained to tunnel handling ^58^ daily during the habituation phase and tunnel handling was used throughout the whole experiment.

At the age of eleven weeks mice were provided with cage enrichment. Cages were cleaned weekly and each mouse was subjected to a visual health check. The enrichment scheme consisted of permanently provided items (running disc with mouse igloo, paper nesting, cotton rolls, Table 2) and five weekly rotating items from structural, housing, nesting and foraging categories (See Table 2 and 3). These enrichment items were randomly exchanged during the weekly cage cleaning. Randomization of the enrichment combination was done with the use of the function randomize() in the software R (version 4.0.4). To motivate the mice in solving the riddles of the foraging enrichment category, a small amount of millet seeds was provided in the morning inside the riddle during the daily animal inspection. Prior to the preference experiments, the mice were used in another experiment (Hobbiesiefken et al. subm. ^45^) but stayed in the above-mentioned housing conditions.

**Table 2:**
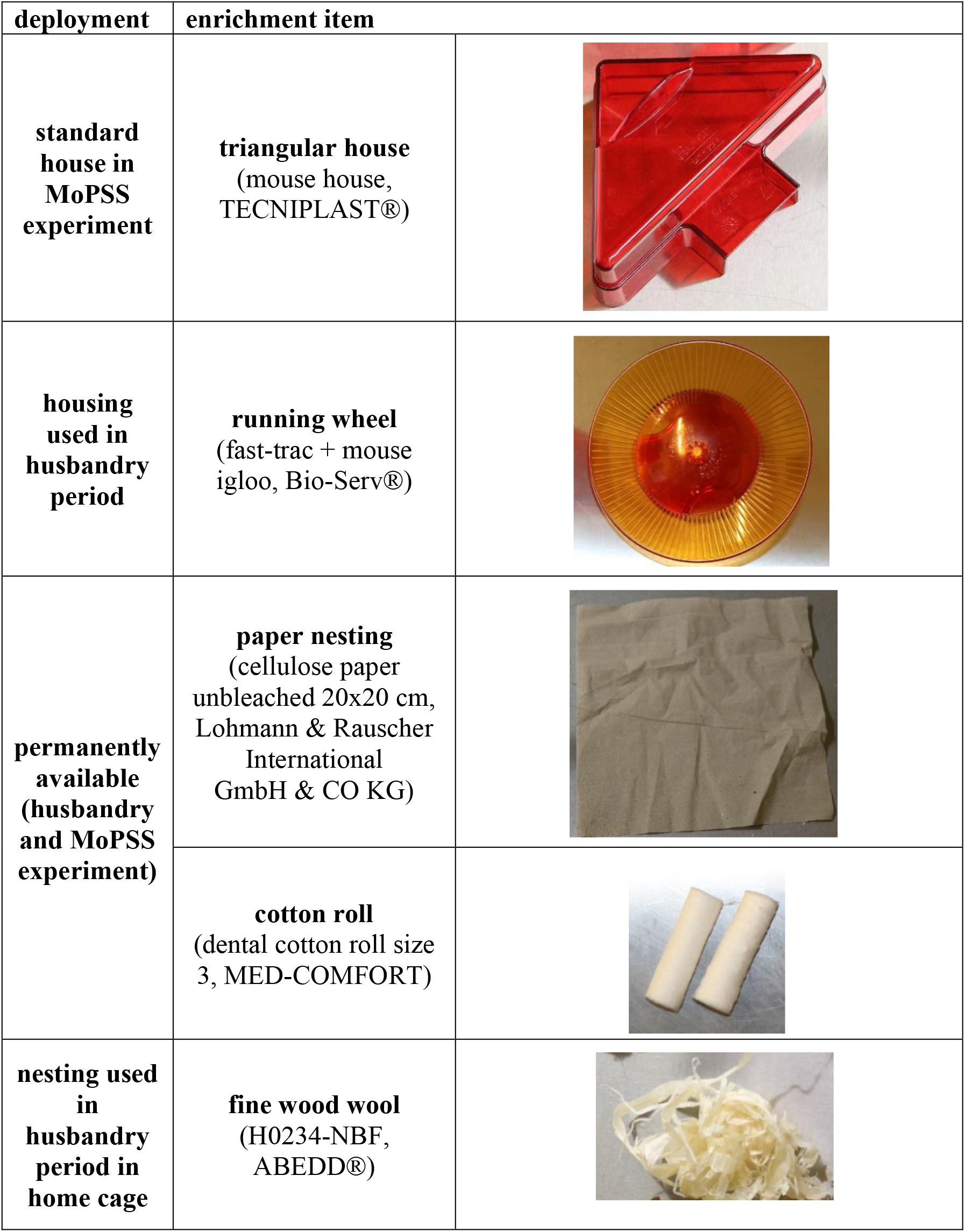
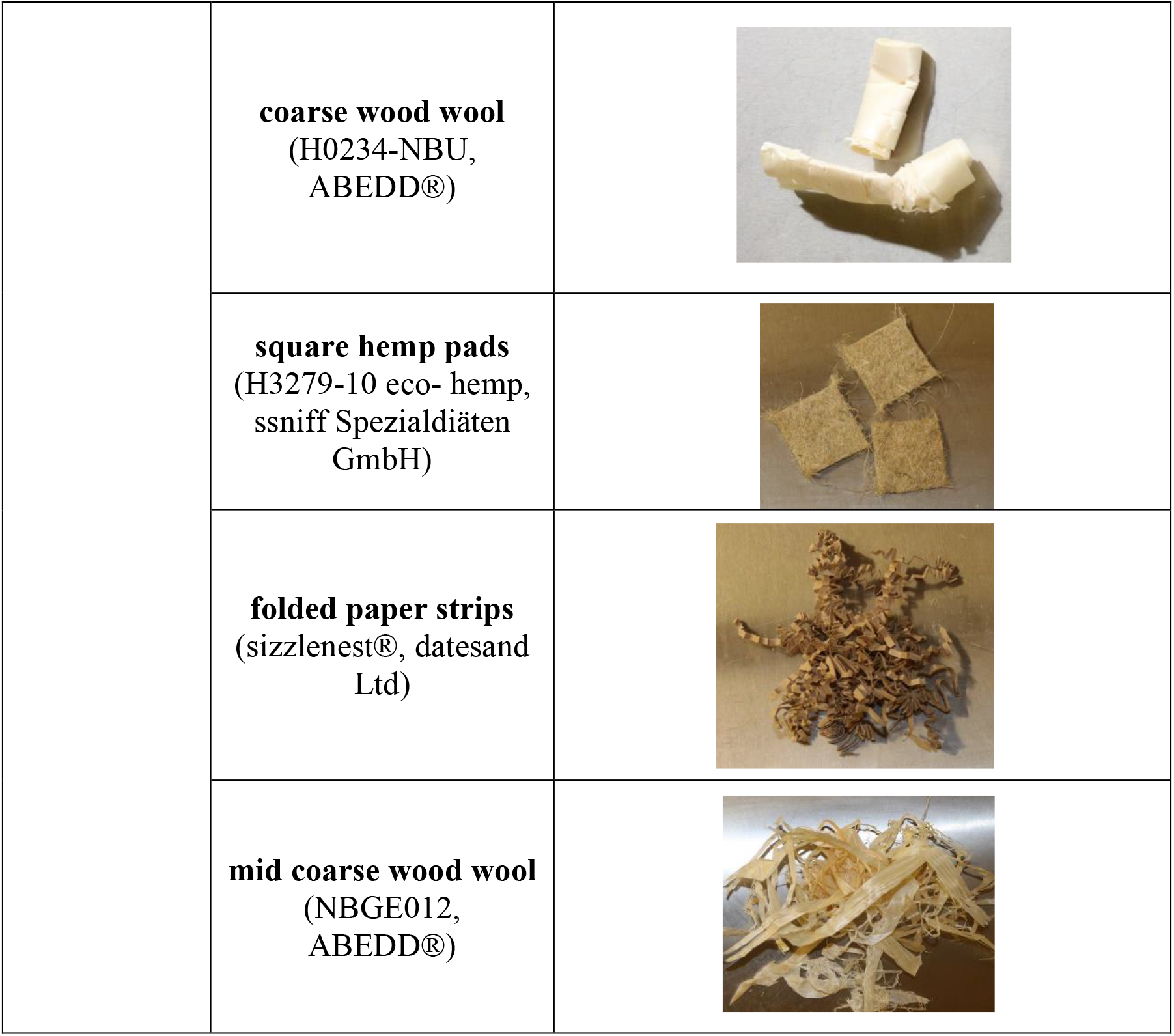
Used enrichment items.

**Table 3.**
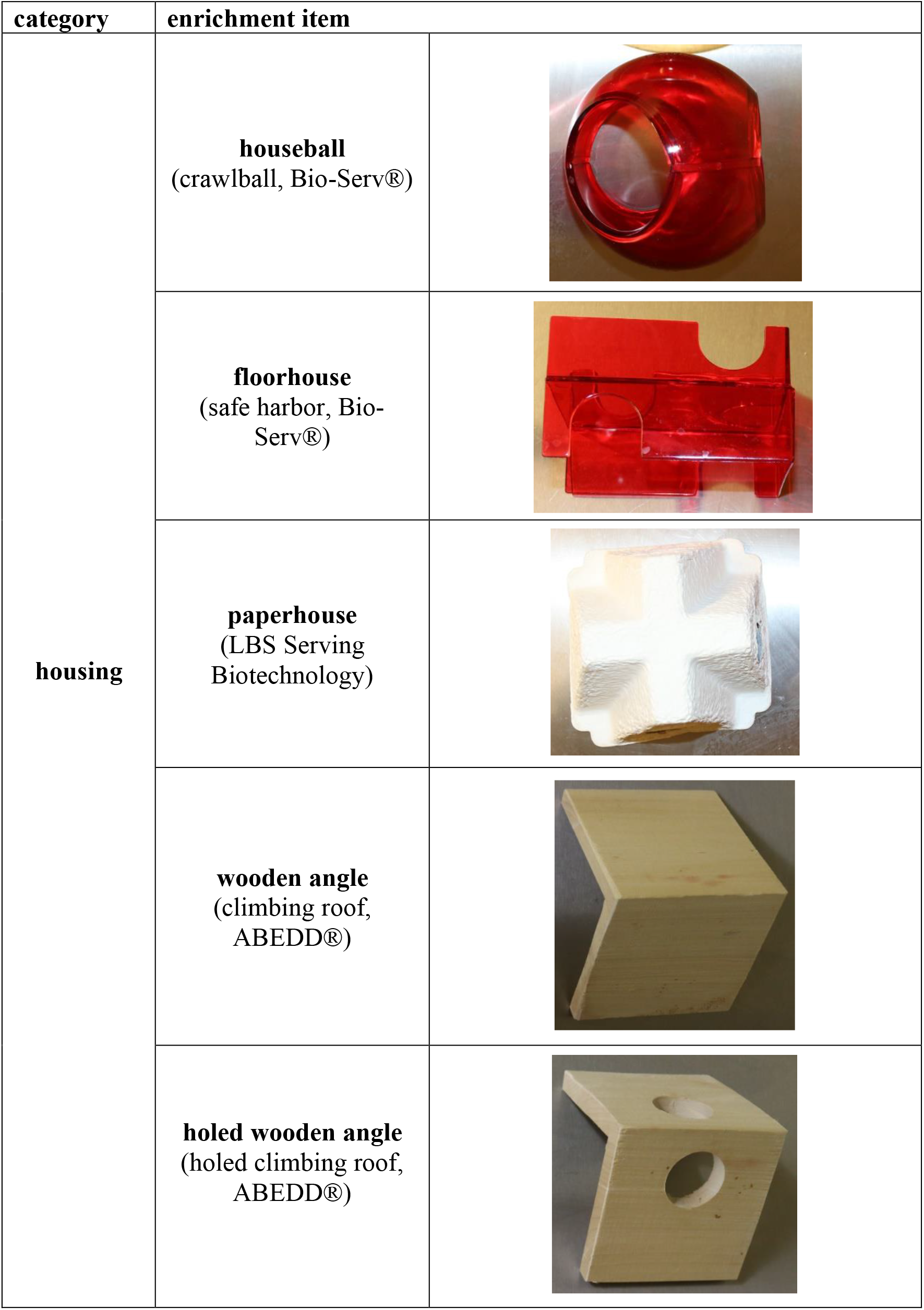
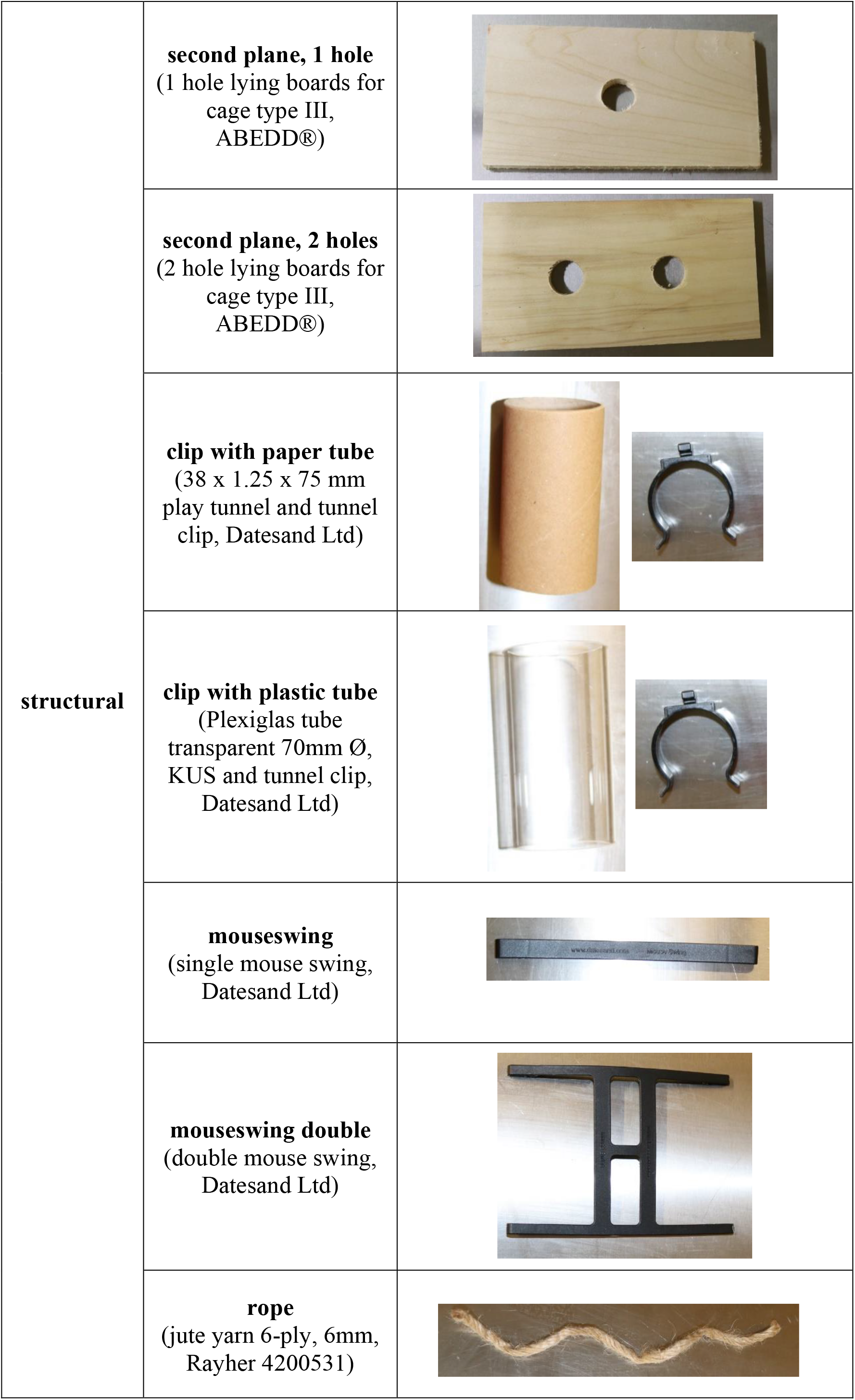
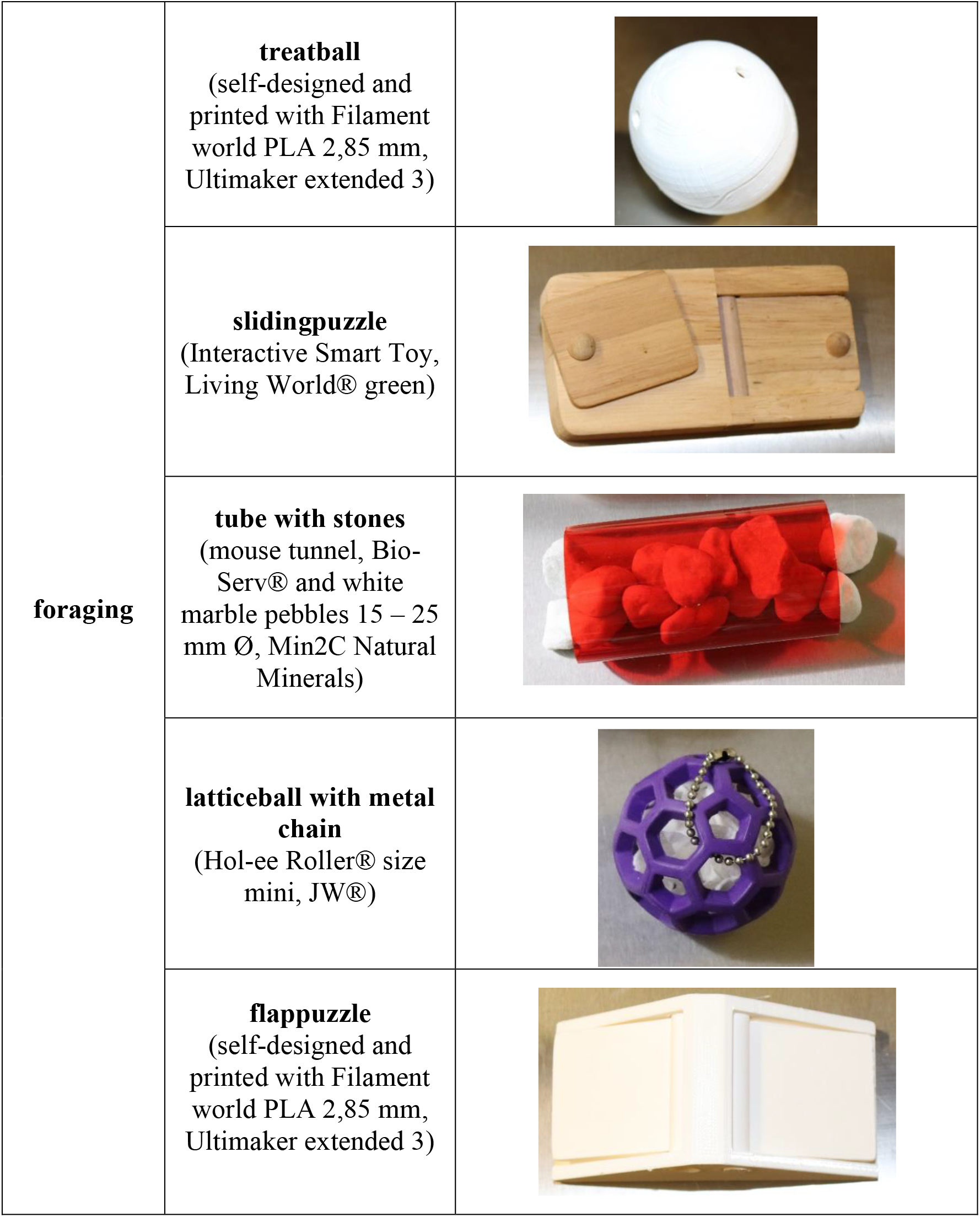
Tested enrichment items.

### Animal identification

For individual animal identification, all animals were provided with a microchip transponder (ISO 11784/85, FDX-B transponders, Planet ID®) under the skin of the dorsal neck region in rostro caudal implantation direction. This procedure took place at the age of 9-10 weeks under general isoflurane anaesthesia and pain reliever (Metacam ®).

Additionally, all mice were color-coded weekly on the tail with a permanent marker (Edding® 750) to distinguish them in video observations.

### Preference testing

After 43 weeks in the enriched housing condition, preference tests were conducted using the Mouse Positioning and Surveillance System (MoPSS) ^31^. The system consisted of two macrolon type III cages, connected with a 30 cm plexiglas tube. Two circular RFID antennas were attached outside the tube. Inside the tube, plastic barriers were installed in order to slow down mouse movement (see Figure 4). The RFID antennas were connected to a reader, which recorded the mouse movements between the left and right cage through detection of the implanted microchip.

**Figure 4.**
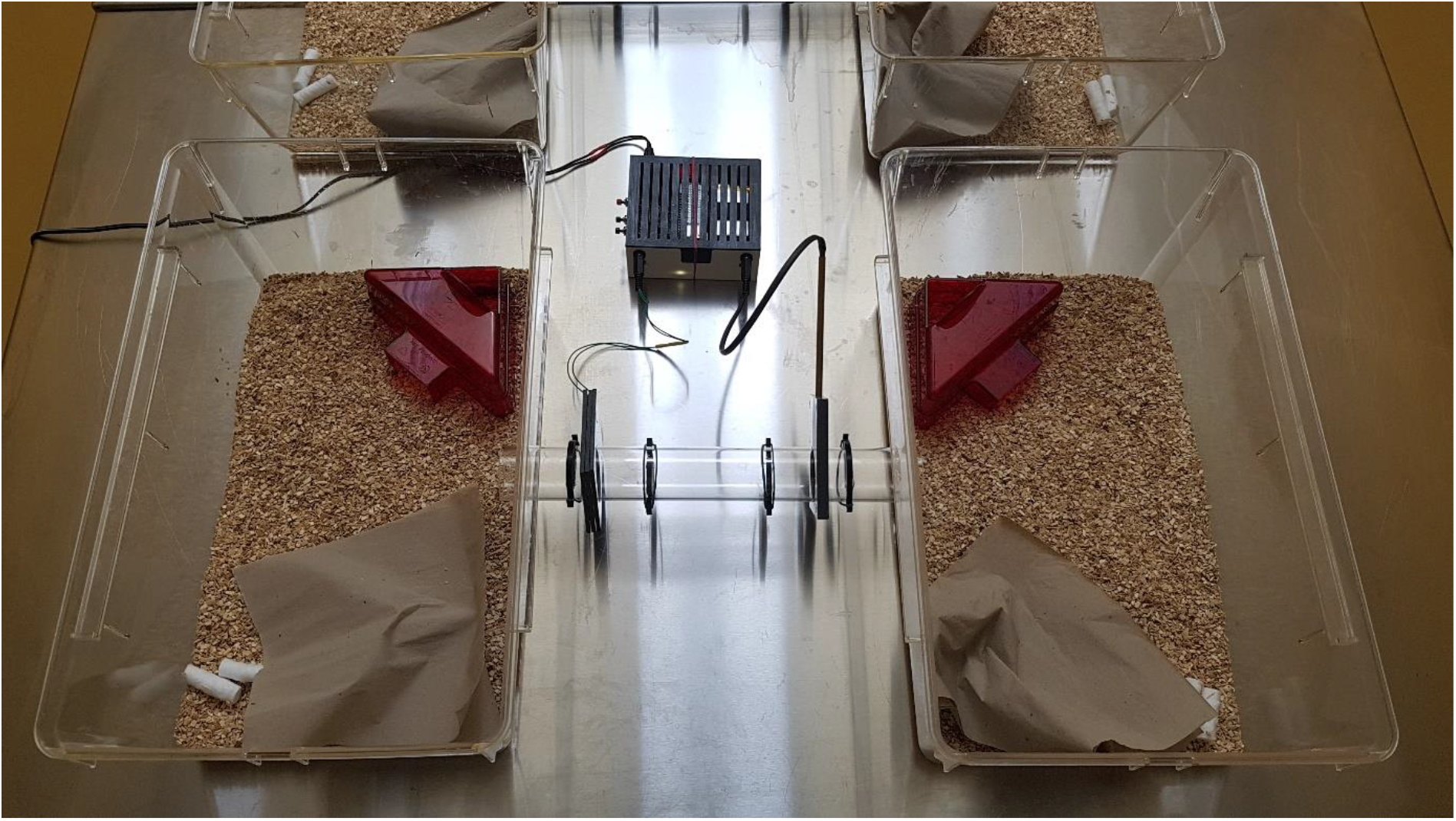
The Mouse Positioning and Surveillance System (MoPSS)

The mice remained in their group of four animals and three preference systems were used in parallel. The systems were positioned in a row on a steel table (see Figure 5: Experimental setup), in an experimental room with the same environmental conditions as during the housing period.

**Figure 5.**
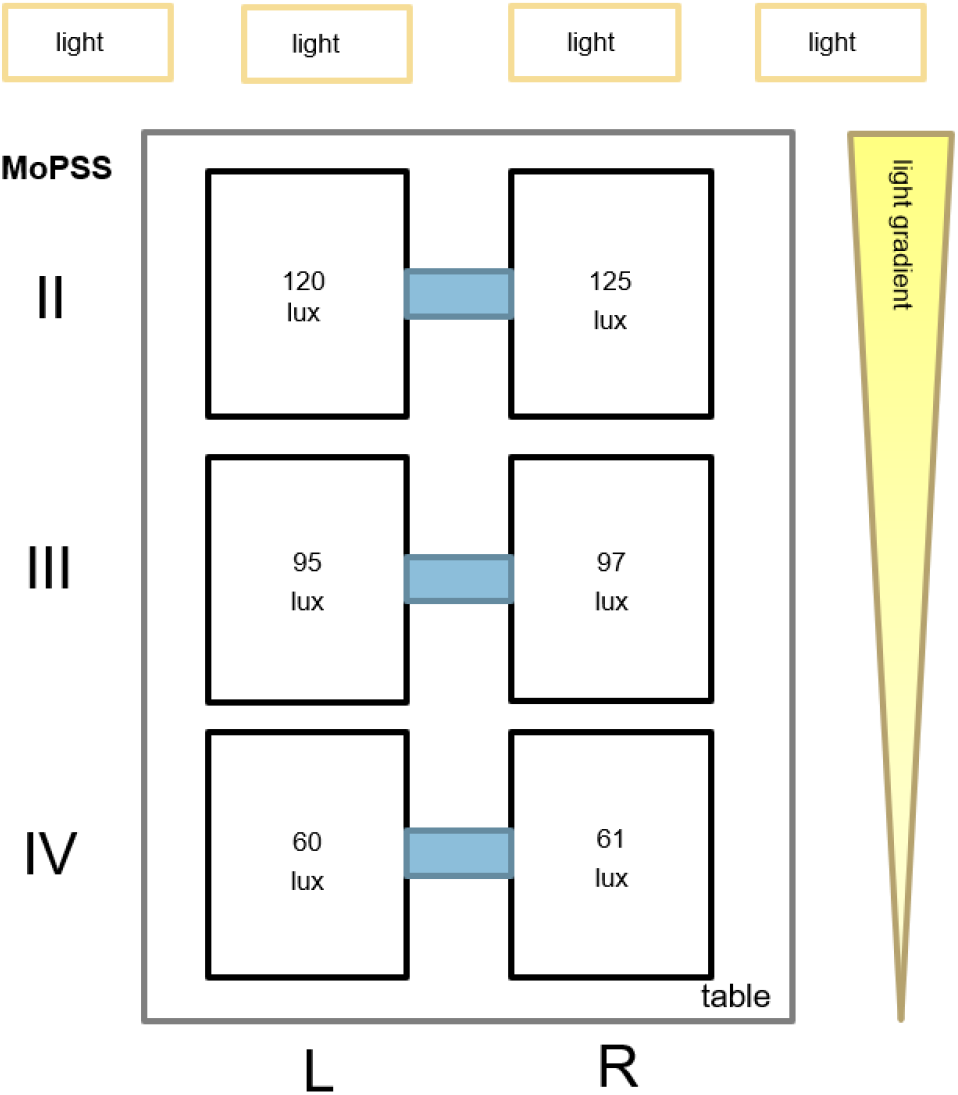
Experimental setup.

To achieve same lightning conditions for the left and right cage of the preference system, four LED lights (Brennenstuhl® Dinora 5000 Baustrahler 47 W 5000 lm Tageslicht-Weiß 1171580) on tripods were set up pointing towards the ceiling. Light intensity in both test cages was checked with a lux meter (voltcraft® light meter MS-1300).

The testing cages were outfitted with 150 g aspen bedding (Polar Granulate 2-3 MM, Altromin), a red translucent triangular plastic house, three uncolored paper towels, two cotton rolls, and water and rodent food (autoclaved pellet diet, LAS QCDiet, Rod 16, Lasvendi, Germany) *ad libitum* with same amount on each side (see Table 2 for equipment details).

Enrichment items placed into the cages were visible so that a full blinded design was not achievable. However, the automated recording of the behavioral data in the absence of the experimenter excluded any possible influence.

Enrichment items of one category each were randomly presented twice for 23 hours starting at 10:00 am until 9:00 am the following day. Between the two sessions using the same items, the enrichment items were switched between the cages to counterballance possible side preferences. In addition also the nesting material and bedding was mixed between the left and right cage and the mice were supplied with their daily amount of millet seeds. The first category tested was the ‘structural enrichments’ followed by the ‘foraging enrichments’ and the ‘housing enrichments’. Two days before the first preference test, the mice were introduced to the experimental setup including the MoPSS for habituation purpose. After completion of the experiments in this work, the animals remained in their housing conditions and were used for further studies.

### Analysis of preference

The mouse tag detections were automatically saved onto a microSD card during the experiment and each detection was marked by a current timestamp with the antenna number (left/right) and the individual mouse RFID tag number. Data analysis and sanity checks with logical correction of missing detections were done using a data evaluation script in the software R (R version 4.0.4, R Studio version 1.3.959) specially developed for MoPSS data analysis ^31^. No missing data were found, all mice were regularly detected and none had to be excluded from analysis of stay times. Stay times for each of the twelve mice in each cage side were calculated as times between cage changes when a mouse tag was detected at a new cage. It has been shown that the time spent in the tube is negligible for preference calculation ^31^ and therefore we did not subtract the time spent in the passaging tube. Stay durations over the 46 hours testing period of each single experiment were summed up for each mouse and then calculated as percentage of the total time. All data was analyzed both at group/cage level and in relation to the length of stay of all individual mice over the total period of 46 hours and over the light and dark phase representing the activity phases of the mice. The calculated percentages of stay durations were then used for comparison of side preferences (left vs. right cage) for enrichment one and enrichment two including a side switch of the presented items. The raw data with stay durations in percentage during the hole 46 hours testing period and divided into the active and inactive time period can be found in the supplementary material (Supplement: Table 1,2,3).

To rank the tested enrichment items regarding the strength of the preference for each item, a method developed by Hatzinger et al. 2012 ^33^ of combining the multiple single binary choices to a ‘worth value’ was performed using R and the package simsalRbim ^34^. A similar method was used by Hopper et al. 2019 to determine the worth value of different items of food rated by a male gorilla ^59^. In short, to estimate the position of an item, the ‘worth value’ of each enrichment item was calculated based on the prefmod package ^33^ with its fit to a log-linear Bradley-Terry model (LLBT). The LLBT was specifically made for paired-comparison testing and estimates a subject’s relative ‘worth value’ for each choice on a preference scale that sums to 1 ^33^. Greater preference is represented here by a higher ‘worth value’.

To determine the agreement amongst the mice regarding the ‘worth value’ for each ranked enrichment item and its estimated position on the scale, a consensus error was also calculated using the simsalRbim package^34^. A detailed example of the calculation of the consensus error can be found on the simsalRbim homepage.^34^ In brief, the consensus error reflects the extent of agreement that the mice showed regarding the preference for a certain enrichment in binary choices over the other tested enrichment items. A value of 0 % points to a perfect agreement of a ranking position and 100 % indicates a full disagreement of all individual mice. It should be noted that CE is biased by the number of individuals, with low numbers resulting in CE being significantly more affected by a single animal. In our presentation of the cage wise preferences we therefore refrained from calculating the CE as the ratings are based on a choice of only 4 animals.

All analyses were run in R version 4.0.4 using RStudio (Version 1.3.959).

### Sample size

It is debated whether or not group housed animals can unequivocally considered to act independently in their choice and therefore each cage would have to be considered as one independent sample ^31,35,36,60^. This presents a dilemma because the mice would either have to be housed individually or the total number of experimental animals would have to be increased by the use of additional cages. As we are explicitly interested in the preference for enrichment items under common social conditions, housing mice singly was not an option. With regard to keeping the overall number of experimental animals as low as possible in the light of the 3Rs, we calculated that 12 mice would be a reasonable sample size if they indeed act independently. In order to demonstrate that individual preference was an independent choice, we conducted a follow and influence behavior analysis using R (Version 4.0.4) with our obtained experimental data from the MoPSS. A *follow event* was defined as a transition of one mouse directly detected within one second after another mouse. The leading mouse detected in this constellation received an *influencer event*. We further calculated a *follow rate* and *influence rate* as follows:

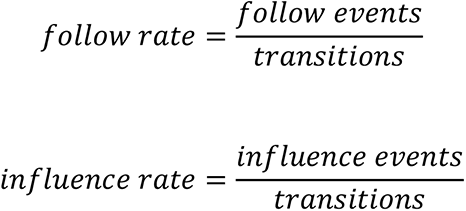

## Acknowledgments

The authors received no specific funding for this work. The work is financed by the annual budget of The German Federal Institute for Risk Assessment (BfR). BfR reports to the Federal Ministry of Food and Agriculture (BMEL). The BMEL had no role in study design, data collection and analysis, decision to publish, or preparation of the manuscript.

The manuscript has been published first on bioRxiv (https://doi.org/10.1101/2021.10.20.465117).

The authors thank the animal caretakers, especially Carola Schwarck and Lisa Gordijenko, for their support in the animal husbandry.

## Authors contribution

All persons who meet authorship criteria are listed as authors, and all authors certify that they have participated sufficiently in the work to take public responsibility for the content, including participation in the concept, design, analysis, writing, or revision of the manuscript.

Conception and design of study: U.H., K.D., L.L.; acquisition of data: U.H.; analysis and/or interpretation of data: U.H., B.U., A.J.

Drafting the manuscript: U.H.; revising the manuscript critically for important intellectual content: K.D., L.L.

Approval of the version of the manuscript to be published: U.H., B.U., A.J., K.D., L.L.

## Supplementary Material

Supplementary Material such as 3D printing templates and raw data tables can be found under: https://github.com/DasDritteR/MoPSS-preference-test-supplements

## References

1. The European Parliament & The Council of the European Union. Directive 2010/63/EU of the European Parliament and of the Council of 22 September 2010 on the protection of animals used for scientific purposes. (2010).

2. Newberry, R. C. Environmental enrichment: Increasing the biological relevance of captive environments. Appl. Anim. Behav. Sci. 44, 229–243 (1995).

3. Olsson, I. A. S. & Dahlborn, K. Improving housing conditions for laboratory mice: A review of ‘environmental enrichment’. Lab. Anim. 36, 243–270 (2002).

4. Lewejohann, L. & Sachser, N. Evaluation of different housing conditions for male laboratory mice by means of preference tests. KTBL SCHRIFT 170–177 (2000).

5. Duncan, I. J. H. & Olsson, I. A. S. Environmental enrichment: from flawed concept to pseudo-science. in Proceedings International Congress of th e ISAE 2001, Davis, USA (2001).

6. Hutchinson, E., Avery, A. & VandeWoude, S. Environmental enrichment for laboratory rodents. ILAR J. 46, 148–161 (2005).

7. Bailoo, J. D. et al. Effects of Cage Enrichment on Behavior, Welfare and Outcome Variability in Female Mice. Front. Behav. Neurosci. 12, (2018).

8. Tang, Y. P., Wang, H., Feng, R., Kyin, M. & Tsien, J. Z. Differential effects of enrichment on learning and memory function in NR2B transgenic mice. Neuropharmacology 41, 779–790 (2001).

9. van Praag, H., Kempermann, G. & Gage, F. H. Neural consequences of enviromental enrichment. Nat. Rev. Neurosci. 1, 191–198 (2000).

10. Kempermann, G., Kuhn, H. G. & Gage, F. H. More hippocampal neurons in adult mice living in an enriched environment. Nature 386, 493–495 (1997).

11. Benaroya-Milshtein, N. et al. Environmental enrichment in mice decreases anxiety, attenuates stress responses and enhances natural killer cell activity. Eur. J. Neurosci. 20, 1341–1347 (2004).

12. Bailoo, J. D. et al. Effects of Cage Enrichment on Behavior, Welfare and Outcome Variability in Female Mice. Front. Behav. Neurosci. 12, (2018).

13. van de Weerd, H. A. et al. Effects of Environmental Enrichment for Mice: Variation in Experimental Results. J. Appl. Anim. Welf. Sci. 5, 125–137 (2002).

14. André, V. et al. Laboratory mouse housing conditions can be improved using common environmental enrichment without compromising data. PLoS Biol. 16, 1–24 (2018).

15. Baumans, V., van Loo, P. L. P. & Pham, T. M. Standardisation of environmental enrichment for laboratory mice and rats: Utilisation, practicality and variation in experimental results. Scand. J. Lab. Anim. Sci. 37, 101–114 (2010).

16. Gross, A. N., Richter, S. H., Engel, A. K. J. & Würbel, H. Cage-induced stereotypies, perseveration and the effects of environmental enrichment in laboratory mice. Behav. Brain Res. 234, 61–68 (2012).

17. Key, D. Environmental Enrichment Options for Laboratory Rats and Mice. Lab Anim. (NY). 33, 39–44 (2004).

18. van de Weerd, H. A. Environmental Enrichment for Laboratory Rodents: Preferences and Consequences. (1996).

19. Baumans, V. Environmental enrichment for laboratory rodents and rabbits: Requirements of rodents, rabbits, and research. ILAR J. 46, 162–170 (2005).

20. Dawkins, M. S. From an animal’s point of view: Motivation, fitness, and animal welfare. Behav. Brain Sci. 13, 1–9 (1990).

21. Baumans, V. & Van Loo, P. L. P. How to improve housing conditions of laboratory animals: The possibilities of environmental refinement. Vet. J. 195, 24–32 (2013).

22. Lewejohann, L., Schwabe, K., Häger, C. & Jirkof, P. Impulse for animal welfare outside the experiment. Lab. Anim. 54, 23677219891754 (2020).

23. Habedank, A., Kahnau, P., Diederich, K. & Lewejohann, L. Severity assessment from an animal’s point of view. Berl. Munch. Tierarztl. Wochenschr. 31, 304–320 (2018).

24. Dawkins, M. S. A user’s guide to animal welfare science. Trends Ecol. Evol. 21, 77–82 (2006).

25. Lewejohann, L. et al. Environmental bias? Effects of housing conditions, laboratory environment and experimenter on behavioral tests. Genes, Brain Behav. 5, 64–72 (2006).

26. Wahlsten, D. et al. Different data from different labs: Lessons from studies of gene-environment interaction. J. Neurobiol. 54, 283–311 (2003).

27. van Loo, P. L. P., Blom, H. J. M., Meijer, M. K. & Baumans, V. Assessment of the use of two commercially available environmental enrichments by laboratory mice by preference testing. Lab. Anim. 39, 58–67 (2005).

28. Sherwin, C. M. Preferences of laboratory mice for characteristics of soiling sites. Anim. Welf. 5, 283–288 (1996).

29. Koolhaas, J. M., van Loo, P. L. P., van Zutphen, L. F. M., van de Weerd, H. A. & Baumans, V. Preferences for nesting material as environmental enrichment for laboratory mice. Lab. Anim. 31, 133–143 (2007).

30. van Loo, P. L. P., Blom, H. J. M., Meijer, M. K. & Baumans, V. Assessment of the use of two commercially available environmental enrichments by laboratory mice by preference testing. Lab. Anim. 39, 58–67 (2005).

31. Habedank, A., Urmersbach, B., Kahnau, P. & Lewejohann, L. O mouse, where art thou? The Mouse Position Surveillance System (MoPSS) - An RFID based tracking system. Behav. Res. Methods 1–23 (2021) doi:10.3758/s13428-021-01593-7.

32. Mekada, K. et al. Genetic differences among C57BL/6 substrains. Exp. Anim. 58, 141–149 (2009).

33. Hatzinger, R. ; R. D. prefmod: An R Package for Modeling Preferences Based on Paired Comparisons, Rankings, or Ratings. Wiley Interdiscip. Rev. Comput. Stat. 1, 128–129 (2012).

34. Talbot, S., Pfefferle, D., Brockhausen, R. & Lewejohann, L. simsalRbim - A package for preference test simulations. https://talbotsr.com/simsalRbim/index.html.

35. Parsons, N. R., Teare, M. D. & Sitch, A. J. Unit of analysis issues in laboratory-based research. Elife 7, 1–25 (2018).

36. Bello, N. M. et al. Short communication: On recognizing the proper experimental unit in animal studies in the dairy sciences. J. Dairy Sci. 99, 8871–8879 (2016).

37. Gaskill, B. N., Rohr, S. A., Pajor, E. A., Lucas, J. R. & Garner, J. P. Some like it hot: Mouse temperature preferences in laboratory housing. Appl. Anim. Behav. Sci. 116, 279–285 (2009).

38. Freymann, J., Tsai, P.-P., Stelzer, H. & Hackbarth, H. The amount of cage bedding preferred by female BALB/c and C57BL/6 mice. Lab Anim. (NY). 44, 17–22 (2015).

39. Van de Weerd, H. A., Baumans, V., Koolhaas, J. M. & Van Zutphen, L. F. M. Nesting material as enrichment in two mouse strains. in Frontiers in Laboratory Animal Science: Joint International Conference of ICLAS, Scand-LAS and FinLAS 119–123 (1996).

40. Van De Weerd, H. A., Van Loo, P. L. P., Van Zutphen, L. F. M., Koolhaas, J. M. & Baumans, V. Preferences for nesting material as environmental enrichment for laboratory mice. Lab. Anim. 31, 133–143 (1997).

41. Roper, T. J. Nesting material as a reinforcer for female mice. Anim. Behav. 21, 733–740 (1973).

42. Van De Weerd, H. A., Van Loo, P. L. P., Van Zutphen, L. F. M., Koolhaas, J. M. & Baumans, V. Strength of preference for nesting material as environmental enrichment for laboratory mice. Appl. Anim. Behav. Sci. 55, 369–382 (1998).

43. Deacon, R. M. J. Assessing nest building in mice. Nat. Protoc. 1, 1117–1119 (2006).

44. Latham, N. & Mason, G. From house mouse to mouse house: the behavioural biology of free-living Mus musculus and its implications in the laboratory. Appl. Anim. Behav. Sci. 86, 261–289 (2004).

45. Hobbiesiefken, U., Mieske, P., Lewejohann, L. & Diederich, K. Evaluation of different types of enrichment - their usage and effect on home cage behavior in female mice. PloS one subm.

46. Leach, M. C., Ambrose, N., Bowell, V. J. & Morton, D. B. The Development of a Novel Form of Mouse Cage Enrichment. J. Appl. Anim. Welf. Sci. 3, 81–91 (2000).

47. Gjendal, K., Sørensen, D. B., Kiersgaard, M. K. & Ottesen, J. L. Hang on: An evaluation of the hemp rope as environmental enrichment in C57BL/6 mice. Anim. Welf. 26, 437–447 (2017).

48. Van de Weerd, H. A., Van Loo, P. L. P., Van Zutphen, L. F. M., Koolhaus, J. M. & Baumans, V. Preferences for nest boxes as environmental enrichment for laboratory mice. Anim. WELFARE-POTTERS BAR- 7, 11–26 (1998).

49. Kirkden, R. D. & Pajor, E. A. Using preference, motivation and aversion tests to ask scientific questions about animals’ feelings. Appl. Anim. Behav. Sci. 100, 29–47 (2006).

50. Dawkins, M. S. Behavioural deprivation: A central problem in animal welfare. Appl. Anim. Behav. Sci. 20, 209–225 (1988).

51. Mason, G. J. & Burn, C. C. Behavioral Restriction. 98–119 (2011).

52. Gross, A. N. M., Engel, A. K. J. & Würbel, H. Simply a nest? Effects of different enrichments on stereotypic and anxiety-related behaviour in mice. Appl. Anim. Behav. Sci. (2011) doi:10.1016/j.applanim.2011.06.020.

53. Meagher, R. K. & Mason, G. J. Environmental Enrichment Reduces Signs of Boredom in Caged Mink. PLoS One 7, (2012).

54. Burn, C. C. Bestial boredom: a biological perspective on animal boredom and suggestions for its scientific investigation. Anim. Behav. 130, 141–151 (2017).

55. Meagher, R. K. Is boredom an animal welfare concern? Anim. Welf. 28, 21–32 (2019).

56. Richter, S. H. & von Kortzfleisch, V. Reply to ‘It is time for an empirically informed paradigm shift in animal research’. Nat. Rev. Neurosci. 21, 661–662 (2020).

57. Russell, W. & Burch, R. The principles of Humane Experimental Technique. (Methuen, 1959).

58. Hurst, J. L. & West, R. S. Taming anxiety in laboratory mice. Nat. Methods 7, 825–826 (2010).

59. Hopper, L. M., Egelkamp, C. L., Fidino, M. & Ross, S. R. An assessment of touchscreens for testing primate food preferences and valuations. Behav. Res. Methods 51, 639–650 (2019).

60. Kappel, S., Hawkins, P. & Mendl, M. T. To Group or Not to Group? Good Practice for Housing Male Laboratory Mice. Animals 7, (2017).

